# Somnus, a tunable, open-source platform for automated EEG sleep scoring in physiological and disease mouse models

**DOI:** 10.64898/2026.10.05.756738

**Authors:** Josh Brenner, Matthew S. Caudill, Huda Y. Zoghbi

## Abstract

Many studies using mouse models of human brain disorders rely on extended recordings of brain electrical activity through electroencephalogram, or EEG. The features that researchers measure in these recordings, such as seizures and epileptiform activity, are deeply influenced by whether the animal is awake, in non-rapid eye movement (NREM) sleep, or in rapid eye movement (REM) sleep. Because total sleep duration varies naturally across animals, rigorous experimental comparisons require classifying the sleep state of every few seconds of a recording. Manual annotation is highly labor-intensive, often requiring hours per subject. While automated scoring tools accelerate this process, they are typically trained exclusively on healthy cohorts and frequently fail when applied to disease models that exhibit atypical or fragmented sleep architecture.

Somnus is an open-source Python package designed to automate rodent sleep staging using EEG, muscle activity (EMG), and optional video-tracking data. It includes a robust base model trained on diverse recordings from six independent laboratories, enabling out-of-the-box classification across various equipment setups without requiring prior manual labeling.

A central feature of Somnus is its capacity for rapid adaptation to pathological sleep structure. The software provides a graphical interface that directs researchers to the specific epochs where the base model is least certain. Users can manually correct these targeted epochs in a built-in viewer by comparing the model’s classification to the raw EEG signal and video, and then use their corrections to fine-tune the model. This workflow empowers researchers to generate accurate custom scorers for complex disease models or novel treatments with just a few minutes of manual correction, bypassing the need to hand-score entire experimental cohorts or develop their own models from scratch.

## STATEMENT OF NEED

Rodent electroencephalography (EEG) remains the gold standard for investigating many neurological disorders, including epilepsy. Crucially, nearly every physiological signal of interest— epileptiform spikes, seizures, and neural oscillations, as well as their responses to pharmacological interventions —is heavily modulated by the animal’s sleep state [Harvey et al., 2025]. Because individual rodents naturally sleep for highly variable amounts of time during a given recording, rigorous experimental design demands state-matched comparisons to isolate true disease phenotypes from state-dependent artifacts. This requirement creates a severe analytical bottleneck. Animal models of neurological disease, such as Rett syndrome [RTT], frequently exhibit both disrupted sleep architecture and abnormal sleep-state signatures [Al Maghribi et al., 2025]. Automated sleep-scoring tools trained predominantly on healthy wild-type (WT) mice frequently fail to generalize to these atypical profiles.

To compensate, researchers are currently forced to choose between manually scoring entire experimental cohorts—a labor-intensive process that is prone to variability between scorers and potential reproducibility concerns—or manually scoring a subset of data and training a new machine learning model from scratch. For laboratories that iterate through multiple distinct genetic lines, pharmacological interventions, or disease paradigms, building a bespoke model for every experimental variant is a prohibitive drain on time and resources.

We designed Somnus for systems neuroscientists and electrophysiologists who require high-throughput automated sleep scoring that can be easily adapted to disease model systems with variable EEG sleep signatures. The base Somnus model is pre-trained on a public dataset featuring diverse recording setups [Rose et al., 2025a; Rose et al., 2025b] and offers strong out-of-the-box performance on wild-type animals. It features an accessible user interface engineered specifically for fast and easy adaptation; Somnus automatically directs researchers to low-certainty epochs that can be manually corrected in-GUI and used to fine-tune the classifier. Researchers can thus generate bespoke classifiers for individual disease model systems with minimal effort, clearing a significant bottleneck in translational EEG-research.

## STATE OF THE FIELD

Automated polysomnography in rodent models has seen great advancement with the introduction of both web platforms and local algorithmic classifiers. Platforms such as Sleepyrat [SleepyRat, n.d.] provide out-of-the-box sleep scoring for EEG/EMG recordings using a closed, subscription-based model. While highly effective for wild-type or standard pharmacological studies, these platforms cannot be inspected, rerun locally, or adapted by users to new phenotypes. They often struggle with the atypical EEG/EMG signatures inherent to neurological disease models—such as the epileptiform spiking, altered spectral power, or highly fragmented sleep architecture seen in RTT and acquired epilepsy paradigms.

On the other end of the spectrum, a variety of open-source, locally executed scoring packages have emerged. AccuSleePy [Barger, 2026; Barger et al., 2019] ships a compact convolutional neural network for standard mouse EEG/EMG, while Somnotate [Brodersen et al., 2024] fits a linear discriminant plus hidden Markov model to a laboratory’s specific annotations and accepts any input modality. These packages address the reproducibility limitations of closed, hosted models. However, these existing tools require the user to train an entirely new model from scratch when neurological disease models fail to conform to established sleep-state paradigms.

We developed Somnus specifically to resolve this methodological gap. Rather than requiring researchers to train a network from the ground up, Somnus treats its shipped model as a robust prior. Corrections made in its review interface update the regression weights by an amount optimized through cross-validation on the laboratory’s own recordings. This ensures that a small set of relabeled epochs moves the model as far as the new evidence warrants and no farther. By anchoring these updates to a strong base model, Somnus provides rapid, stable adaptation to atypical disease phenotypes that can be established on a small amount of data.

## SOFTWARE DESIGN AND METHODS

Somnus is organized as a five-stage workflow (Figure 1): recordings are loaded and per-epoch features computed, a logistic classifier scores each epoch, the least-certain epochs are flagged for review, the user corrects them, and these corrections are used to fine-tune the model. EEG spectral and EMG features are computed on 4-second epochs, using Openseize for spectral estimation [Caudill, 2023], and z-scored within each recording. This lets a single model accept recordings from different amplifiers, pre-processing pipelines, and electrode setups without per-subject calibration, at the cost of requiring each recording to contain both sleep and wake. Features that a recording cannot measure are set to zero, the mean after standardization, so they neither favor nor oppose any state.

**Figure 1.**
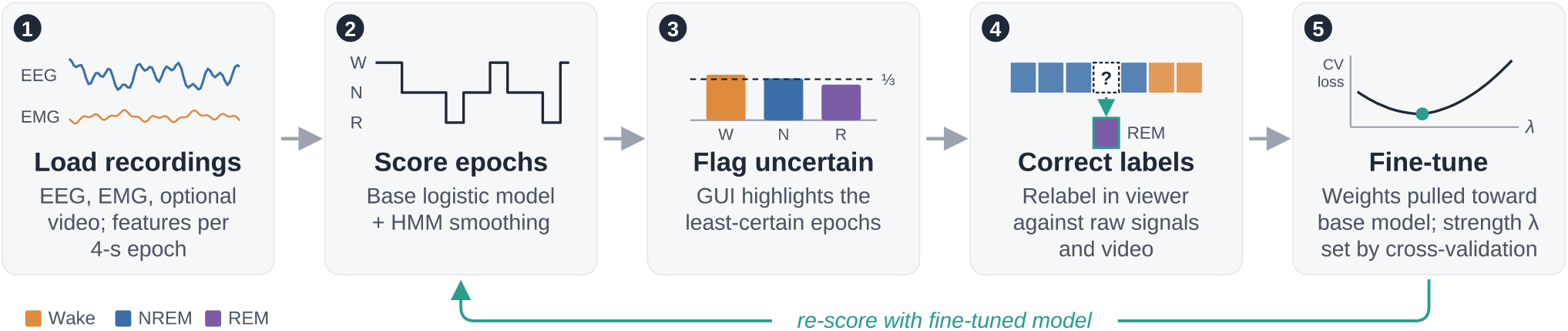
Somnus workflow. (1) Recordings are read in and per-epoch EEG/EMG (and optional video) features are computed. (2) The base model scores each epoch, followed by HMM smoothing. (3) The GUI flags epochs whose class probabilities are near chance. (4) The user corrects these labels in a viewer showing the raw signals and video. (5) The logistic weights are fine-tuned with a penalty toward the base model, with penalty strength λ chosen by cross-validation, and the recordings are re-scored.

The classifier is a logistic model. Its small parameter count keeps fine-tuning on small label sets stable, and its weights remain interpretable. Temporal structure is handled separately by a hidden Markov model with Viterbi decoding [Viterbi, 1967], whose transitions can be adjusted manually or re-estimated algorithmically from labeled data. Manual adjustment raises the transition matrix to a single “stickiness” exponent, while re-estimation can be set to either add to or replace observed transition counts to counts from the base model.

Fine-tuning minimizes class-balanced loss on the user’s labels, with a penalty in favor of the shipped model’s weights. Updates to the coefficients of the logistic model can be found by minimizing the L2-penalized log-linear function L(w) + λ‖w − w_0_‖^2^, where w_0_ are the shipped model’s coefficients, for each value of the penalty parameter λ. This parameter is selected by cross-validation by this software and controls how strongly new data changes the coefficients. Candidate strengths are compared against the unmodified model under leave-one-recording-out validation.

## RESULTS AND RESEARCH IMPACT

We compared Somnus and SleepyRat against expert hand-scoring, reporting Cohen’s κ [Cohen, 1960] both overall and for individual sleep states. On wild-type recordings, both models agreed closely with expert scoring (overall κ = 0.98 for SleepyRat, 0.96 for Somnus). To test their performance in a disease model, we next scored recordings from mice with Rett syndrome (RTT). RTT models present a unique scoring challenge, as they feature highly fragmented sleep architecture and abnormal EEG/EMG signatures. REM sleep in RTT mice is particularly difficult to score due to the frequent presence of epileptiform activity. Consequently, when tested on RTT recordings, the performance of both baseline models degraded significantly (overall κ = 0.52 and 0.66), and neither model could meaningfully detect REM sleep (REM κ = 0.12 and −0.02, respectively).

Fine-tuning enabled Somnus to rapidly adapt to these atypical signatures with minimal user input (Figure 2). Manually rescoring just 13 minutes of hand-labeled RTT recordings restored much of its performance, raising the overall κ to 0.74 (NREM 0.76, REM 0.23, Wake 0.79). Expanding this fine-tuning set to 2.2 hours of labels further improved the overall κ to 0.80 (Wake 0.84, NREM 0.82, REM 0.32). While REM remains the hardest state to classify—reflecting the severely disrupted, seizure-laden REM of RTT mice—these results demonstrate that relatively little retraining can recover highly usable scoring on a complex disease phenotype where rigid, out-of-the-box tools fail outright.

**Figure 2.**
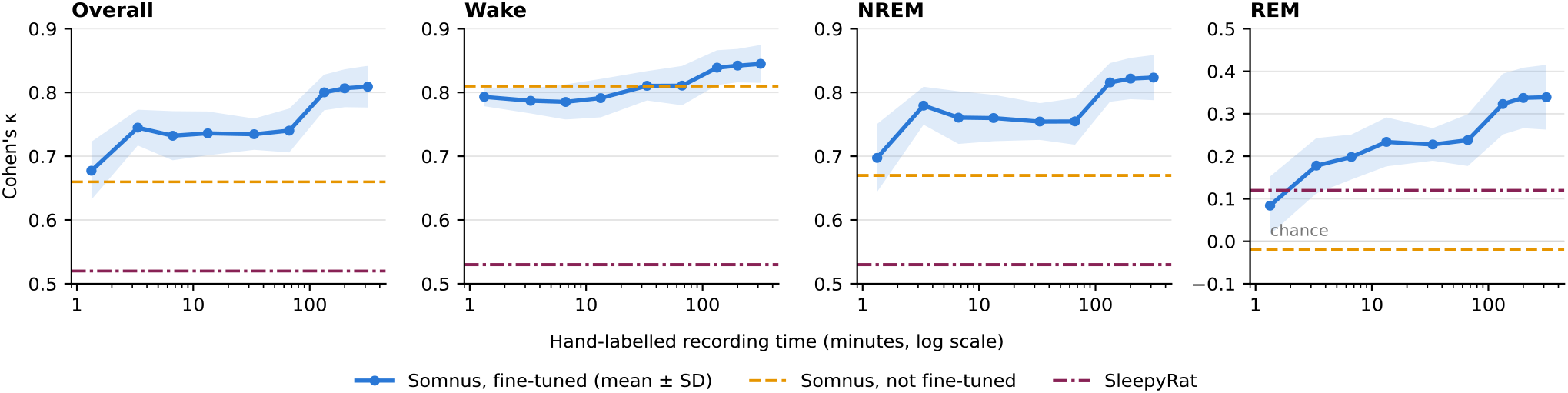
Cohen’s κ against hand-scoring on held-out epochs from RTT mouse recordings, overall and for each state. Blue: Somnus fine-tuned on increasing amounts of hand-labelled recording (mean ± SD, 10 random splits). Dashed lines: Somnus without fine-tuning (orange) and SleepyRat (pink).

## AI USAGE DISCLOSURE

Generative AI was used during software development. Claude Fable 5.1 and Opus 5.5 (Anthropic) and Gemini Pro 3.1 (Google) assisted with writing code and structuring and editing the manuscript under the author’s direction. The scientific questions, model design, feature engineering, fine-tuning approach and evaluation strategy were conceived by the author. The authors reviewed, tested and validated all AI-assisted code, and takes full responsibility for the correctness and integrity of the software and this paper.

## CODE AVAILABILITY

Somnus is available at https://github.com/joshbrenner/somnus and can be installed from PyPI (`pip install somnus-eeg`) under the BSD 3-Clause license.

## ACKNOWLEDGEMENTS

This work was supported by a National Institute of Neurological Disorders and Stroke (NINDS) Ruth L. Kirschstein National Research Service Award (F32NS142584; J. Brenner, PI), NINDS grants 2R37NS057819-21 (H.Y. Zoghbi, PI) and 2R01NS100738-A1 (J. Tang, PI; M.S. Caudill, Co-PI), and the Howard Hughes Medical Institute (H.Y. Zoghbi, Investigator). We thank Claudi Singhai for help with sleep scoring, and Rose et al. and the five contributing laboratories for sharing the Mouse Sleep Staging Validation dataset (MSSV; OpenNeuro ds006366; Rose et al., 2025a), which made training the base model possible.

## Notes

https://github.com/joshbrenner/somnus

https://pypi.org/project/somnus-eeg/

### Competing Interest Statement

The authors have declared no competing interest.

